# Australian researchers’ perceptions and experiences with stem cell registration

**DOI:** 10.1101/2024.03.11.584334

**Authors:** Mengqi Hu, Dan Santos, Edilene Lopes, Dianne Nicol, Andreas Kurtz, Nancy Mah, Sabine C. Muller, Rachel A. Ankeny, Christine A. Wells

## Abstract

The recently issued ISSCR standards in stem cell research recommend registration of human pluripotent stem cell lines (hPSCs). Registration is an important part of establishing stem cell provenance and connecting cell lines to data derived on those lines. In this study, we sought to understand common barriers to registration, by conducting interviews with forty-eight Australian stem cell stakeholders, including researchers, clinicians, and industry professionals. Australian stem cell researchers do not routinely register their lines, and of those Australian lines captured by an international registry, only a third have completed the registration process. Most registered Australian cell lines miss information about their ethical provenance or key pluripotency characteristics. Incomplete registration is poorly aligned with the goals of open science that registries are founded on, and users themselves expressed concerns about the quality of the partial information provided to the resource. Registration was considered a publication hurdle, and this impacted on user perceptions of usefulness of registration, and lowered the likelihood that they would engage with registries to find resources. Although the Australian community represents a small fraction of registry users, the results of this study may suggest ways for journals, registries, and the stem cell community to improve registration compliance.

**Highlights:** Researchers who perceive registration as a publication hurdle are unlikely to complete the registration process.

Incomplete registration promotes misunderstandings about the goals and quality of a registry. Full registration of pluripotent stem cell lines is an important step in verifying cell line provenance.

Greater public awareness of registration, combined with requirements for full registration by journals will support the community adopt ISSCR recommendations to register lines.

## 1. Introduction

### 1.1 Registration of stem cell lines facilitates open science

To integrate FAIR principles (Findable, Accessible, Interoperable and Reusable) into stem cell field, the recently published ISSCR Standards for Human Stem Cell Use in Research Guidance 2023 (Ludwig et al., 2023) advocated for the practice of registration of stem cell lines through international resources such as the Human Pluripotent Stem Cell Registry (hPSCreg) (Seltmann et al., 2016). hPSCreg was initiated to inform the European Commission (EC) of the state of human embryonic stem cells (hESCs) research in Europe, and was extended to include induced pluripotent stem cells (iPSCs) soon after that technology was adopted (Kurtz et al., 2022). It works closely with European funders and with research alliances including the international stem cell banking initiative, which creates awareness and uptake of the resource.

Registration of pluripotent stem cell lines is a solution to the poor naming conventions and ambiguous data provenance that has plagued the stem cell field. A naming convention was proposed by Luong and colleagues (2011), who argued that the linking of quality assurance data associated with individual cell lines was confounded by inconsistent naming conventions, a problem exacerbated as the field expanded and more laboratories were generating hPSCs. Ambiguities arise when different lines are assigned identical names (Kurtz et al., 2018), or when multiple synonyms are used for the same line (Luong et al., 2011). The unique identifier (UI) generated by hPSCreg serves as a persistent, unique, and stable accession number on the registry, which can be used to cross-reference synonyms for individual cell lines across different data resources, such as BioSample (Sayers et al., 2022), Cellosaurus (Bairoch, 2018), and Wikidata (Page, 2022). Journals including *Stem Cell Research* have established a publication prerequisite for its Lab Resources articles, necessitating authors to obtain hPSCreg UI for newly described cell lines (*Stem Cell Research: Lab Resources*, 2023). Cell lines with a hPSCreg UI are linked with publications that cite this identifier, allowing users to find data and use-cases associated with that line.

### 1.2 Most pluripotent stem cell lines are not registered

Despite the establishment of registries like hPSCreg, and a decade of journals calling for adoption of standardised cell line nomenclature, the available evidence shows that most hPSCs are not registered, and poor naming practises continues to plague the field. There were at least 12,168 hPSCs publicly known in 2016, which included more than 2,168 hESCs and 10,000 hiPSCs (Guhr et al., 2018), a number that is no doubt an underestimate given the growth of the field. The current status of hPSCreg as of March 2024 was 6,539 registered hPSCreg lines (967 hESCs, 5572 hiPSCs) (hPSCreg, 2024b), which accounts for only 54% of the total number of lines estimated eight years ago. In assessing the effectiveness of the recommendation by ISSCR for registration of hPSCs, it is important to understand why the stem cell community are not currently adopting the practice of registration.

### 1.3 Australia lacks historical drivers that have supported registration elsewhere

Australia was an early adopter of hPSC research, and a series of significant national funding efforts helped promote and develop the stem cell community in Australia (Finkel, 2008, 2011). As a result, the Australian stem cell community is relatively highly engaged network. Australian funders have adopted open access requirements for publications and data sharing but have not required registration of any cell lines (Australian Research Council, 2021; National Health and Medical Research Council, 2022). In a series of public reviews about the generation and use of embryos, including hESC technologies, the establishment of a national stem cell bank was discussed, but not enacted (Legislation Review Committee, 2005). Prior to this, Australians were expected to use international infrastructure such as the UK Stem Cell Bank (UK Stem Cell Bank, 2024) and hPSCreg. However, only 302 Australian lines are so far registered in the hPSCreg (hPSCreg, 2024a), and three deposited in the UK Stem Cell Bank (UK Stem Cell Bank Repository Statistics, 2021). Australia is therefore a useful testbed to investigate what motivates or impedes the registration of hPSC lines. This article focuses on reporting on the findings from forty eight Australian stem cell researchers, manufacturers and industry professionals who use hPSC, including eight interviewees who had previously registered lines with hPSCreg. Although sampling only a small cross section of the global stem cell community, the themes emerging from this study may nevertheless help the stem cell community understand some of the external drivers for registration, and further sheds light on how user experiences with registries impact on likelihood of further engagement with the stem cell resources held by global registries.

## 2. Material and Methods

### 2.1 Ethics

The ethics approval to undertake this research was granted by the University of Adelaide’s Human Research Ethics Committee (Approval number: H-2022-095). This research was part of a larger study of Enabling Openness in Australian Stem Cell Research (EOAR). The consent form and the participant information sheet are provided in Supplementary Data 1 and Supplementary Data 2 separately. In accordance with ethics obligations, the identities of the interviewees will remain confidential.

### 2.2 Participants

We employed multiple methods to identify potential interviewees. This involved reviewing contacts listed for registered Australian lines in hPSCreg (https://hpscreg.eu/) and Stemcell Knowledge & Information Protal (SKIP, https://saiseiiryo.jp/skip_archive/) registry, collaborating with hPSCreg team to identify potential participants from Australian institutions who had previously registered their lines in hPSCreg, and seeking referrals from existing participants. To identified research labs, we reviewed public funding documents at the Australian Research Council (ARC), National Health and Medical Research Council (NHMRC) and Medical Research Future Fund (MRFF) Stem Cell Therapies mission. We also leveraged previously funded national programs such as the Stem Cells Australia initiative, and networks such as the Australasian Society for Stem Cell Research (ASSCR) to identify the members in the Australian stem cell community. Participation in the project was promoted at the ASSCR Conference 2022 and Lorne Genome 2023. Additionally, some early interviewees suggested new participants through their networks. In total, 61 potential interviewees were contacted via email, with 56 responding, resulting in a 92% response rate. Among these people who responded, eight individuals did not proceed with an interview for various reasons: four stopped responding after an initial expression of interest, two relocated, two directed us to other people. In total, 48 participants were recruited. Recruitment to the study was halted once new themes stopped emerging from the interviews.

All 48 participants (29 males and 19 females) are stem cell stakeholders with (possibly dual) roles including academic researchers (47), clinicians (2), and industry professionals (4). Participants ranged from various career stages, including research group leaders (38), and postdocs (9), spanning age ranges from 25-34 to 65+ (Table 1). Eight participants reported prior experience with hPSCreg.

**Table 1.**
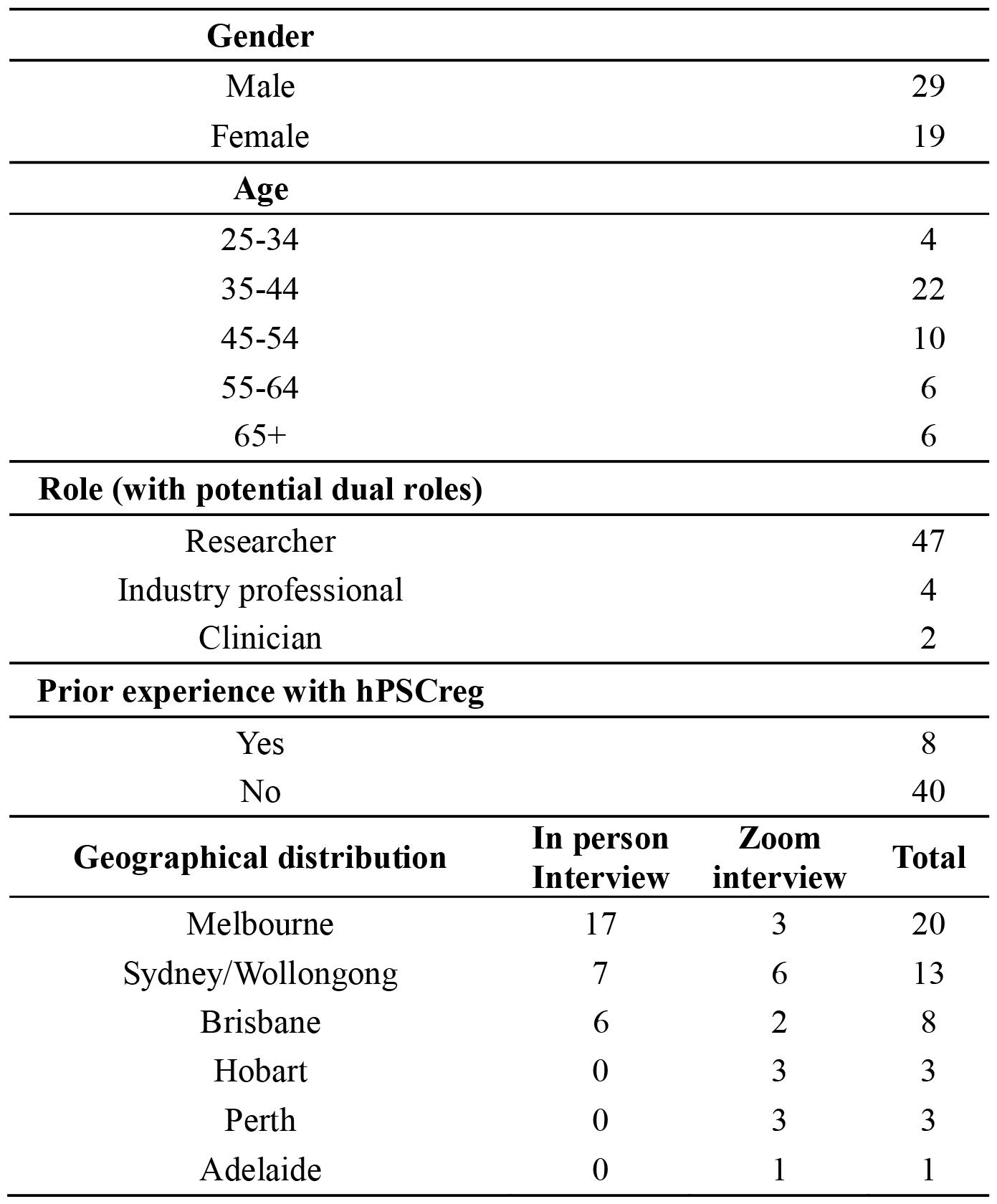
Demographic characteristics of interviewees (n = 48)

### 2.3 Procedure

Initial emails to potential interviewees were sent with an introduction to this project. Once interest in participation was expressed, consent to audio/video-record was obtained prior to the interview. Online survey was provided for quantitative assessment of experience with registries. Interviews were conducted in person or online based on feasibility and interviewee schedules.

Semi-structured interviews were conducted between July 2022 and April 2023. Thirty interviews were conducted in person in Melbourne, Sydney/Wollongong and Brisbane, and eighteen interviews were conducted by video conferencing with participants based in Melbourne, Sydney/Wollongong, Brisbane, Hobart, Perth and Adelaide (Table 1).

### 2.4 Data Collection and Assessment

The interviews focused on delving into participants’ experiences with the registries that they had used. For example, participants were asked to reflect on their motivations for using a registry, describe experiences, and expand on the reasons for the quantitative assessments given in the survey. One participant provided new information through emails post-interview and consented for this piece of information to be treated in the same way as the interview data. Two semi-structured interview guides were used for people with prior experience with a registry and for those without. Interviews ranged from 30 to 90 minutes. Interviews were transcribed verbatim using transcription service Otter and verified by author DS. Following each interview, the interviewers wrote memos with a summary, observations, ideas for future interviews, and potential themes. Five transcripts representing interviewees across different career stages and roles were selected for initial thematic analysis. Author MH created a codebook to capture themes related to registration practises that were identified in the interviews and throughout the research process. The codebook was revised several times throughout the coding process as more themes were identified, and a final version was used to code all the interview transcripts. Interviews were coded with the assistance of NVivo-12. In the following section, interview quotes will be included and analysed; each quote will be identified with a code (e.g. RS01 = Researcher 1) to maintain participant confidentiality.

The purpose of the survey was to capture information about users experiences with registries, using a Likert scale between 1-5 (with 1 = Strongly Disagree, and 5 = Strongly Agree). Answers were used to prompt discussion during the following interview. In total, 21 participants completed the survey, 8 who had used a registry previously, and 13 who had not. The full survey is provided in Supplemental Data 3.

## 3 Results

### 3.1 Few Australian researchers have experienced using a registry

We asked all 48 participants whether they had used a global registry previously, and the majority had not. Our data revealed that most researchers were unfamiliar with international registries, with many confused about what the term ‘registry’ referred to. In the initial three interviews, participants mistakenly assumed that a registry was a biobank repository. To avoid confusion in subsequent interviews, we provided definitions of a registry and a biobank repository. Nevertheless, terms associated with biobanking, such as “deposit” and “shipping”, were used interchangeably when interviewees refered to registry-related topics.

In total, out of 48 participants, only nine reported having used a registry, with the majority (8/9) utilizing hPSCreg, while one mentioned the NIH Embryonic Stem Cell Registry (NIH hESC Registry). Our review of global databases further confirmed that hPSCreg is the most comprehensive catalogue of Australian hPSCs, with 302 Australian hPSCs registered as of 2024 (hPSCreg, 2024). Alongside asking about participants’ experience with a registry, we also enquired about how frequently they generated new hPSC lines. Based on these anecdotal data, we estimate that at least 1,300 hPSCs have been generated by the cross-section of the community that we interviewed. This suggests that only around 23% of Australian hPSCs have been registered on hPSCreg.

### 3.2 Publication is the major motivation for registration

Nine Australian registrants were identified during the recruitment process, and eight of them participated in this study. These users generally agreed that their primary motivation to register was a journal requirement prior to publication. The five respondents interviewed in 2022 stated that they registered to obtain hPSCreg UI based on the standard stem cell nomenclature, as required by the journal *Stem Cell Research* (as exemplified by RS21). However, two of the three users interviewed in 2023, RS34 and RS38, indicated that the updated requirements in *Stem Cell Research* prompted them to complete the entire registration process (full registration). As shown in Figure 1, obtaining the UI involves entering minimum information about the generating institution, cell types (whether iPSC or ESC), and related cell lines; whereas full registration requires completing additional mandatory fields, such as detailing the ethical provenance, derivation, characterization of the cell lines, as well as undergoing a verification process by hPSCreg curators.

**Figure 1.**
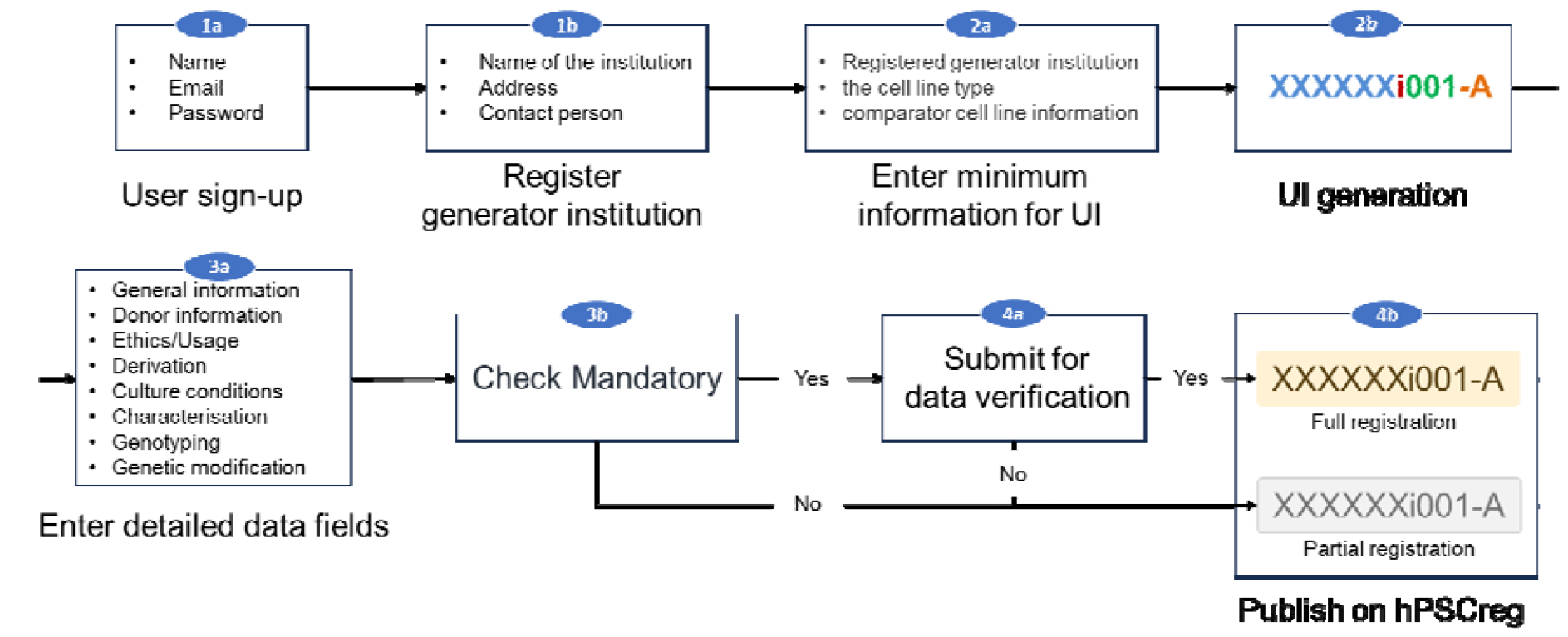
The workflow to obtain a UI or fully register a hPSC line in hPSCreg. To access cell line registration, users must first sign up for an account (1a) and ensure the associated institution is registered (1b). Step 2a allows users to generate a UI (2b). The process can halt at this point and the UI is retained for the cell line from the associated institution. Full registration (step 3a-4b) requires additional mandatory information necessary to establish the ethical and technical provenance of a line. Additional optional information (not shown) is supported for users wishing to share details of additional phenotypic characteristics of a line.

> *“There’s a journal called Stem Cell Research. Stem Cell Research has a resource publication section, which is very formulaic with a number of very specific things you have to do… And one of the very specific formula points is the lines had to be registered with hPSCreg. So that is my very specific reason and sole reason for being involved in any stem cell registry.” (RS21)*
>
> *“I think from memory, the partially registered ones, like the journal requirements only needed there to be a number and then there only needed to be some of those details filled out in the registry. And then, so if that was all that was required, that was all that I did. And then the journal updated their requirements. And so the last publication that I did, I had to fully register.” (RS34)*

### 3.3 Two-thirds of Australian hPSCreg registrations are incomplete

According to hPSCreg website (Figure 1, panel 4b), two-thirds (200/302) of Australian lines remain partially registered, lacking essential details regarding their ethical or technical provenance (Figure 1). All eight Australian hPSCreg registrants had multiple lines published on hPSCreg. While four registrants completed the full registration process for some lines, all had lines that are only partially registered. In response to questions about partial registration, three interviewees explained that they stopped after the UI was obtained, three further expressed an intent to finalise registration later. Respondents were aware that partially registering a cell line may limit their visibility (RS38). Interestingly, two of the research groups who did not register their lines nonetheless recognized the advantages of a UI, adopting hPSCreg nomenclature and its standardized information documentation for internal record-keeping.

> *“Again, part of that was, for example, the lines that are submitted, I feel like they would have very little visibility, especially since I didn’t add some of the specifics about us (our lines). And then they wouldn’t really come up in a search unless you search for that specific line (UIs), which you would do based on the fact that they are published.” (RS38)*

Interviews also reflected limited exploration of the registry itself. Half (4/8) of the registry users reported using hPSCreg to search for cell lines, including two who did so in preparation for the interview. Although hPSCreg includes catalogues of research projects, and jurisdictional legislation and policy in stem cell research, none of the interviewees reported utilizing these functions.

### 3.4 Australian registrants exhibit skepticism regarding the reliability and utility of hPSCreg

Seven hPSCreg registrants expressed varying degrees of negative opinions about the utility of hPSCreg and its registration process. Among them, five generally acknowledged its usefulness but encountered difficulties using the site, while an additional two registrants described the registration process as extremely difficult. After thematic analysis, we observed that while the specifics of negative experiences varied, many of them seemed to be related to misunderstandings about what information was collected by the registry and how hPSCreg validates this data. Two interviewees, RS07 and RS34, expressed concerns about hPSCreg’s request for donor consent forms, citing potential privacy issues for donors and further worrying that these requests might breach their own ethical obligations in deriving cell lines. However, according to hPSCreg documentation, the registry only requires blank or redacted consent forms with no identifiable donor details included. A third respondent, RS34 voiced frustration with the full registration process, but spoke about optional fields as if these were mandatory. In contrast, one researcher who fully understood which fields were mandatory and which were optional, found the data entry process straightforward. RS38 expressed surprise about the rapid verification speed of hPSCreg, and this also prompted questions about how the data was reviewed. There were questions of trust about the quality of information provided by users (RS34) that reflected user-scepticism in the registration process.

“Yes at some point I just feel like they’re asking too much like information for the cell line. Like we have a process to get consent and ethics so sometimes we are not comfortable to share the information of the patient. So that’s why. But it is our first interaction with the registry so I’m still exploring it, what information I should give and what I should not. (RS07)”

> *“And so the last publication that I did, I had to fully register. And it was a little bit more difficult to do the full registry, because, for example, it wanted me to upload ethics, consent forms and things like that, which I have, but I felt like that was a bit of a breach of privacy, because they’ve got people’s names and signatures and addresses and stuff like that on them. And I didn’t want to upload that information.” (RS34)*
>
> *“So one of the things that you do is a trilineage differentiation to make sure it makes ectoderm, endoderm, mesoderm, right, but I stopped there. Like, I’m not gonna go and make muscle cells or a heart or a liver or, you know, I’m just like, okay, it can poke its nose in that direction… it’s too much work to do that and the registry did seem to want a lot of information that I either couldn’t provide or didn’t have time or resources to provide. And so in that sense, it was a bit like not applicable, not applicable, not applicable, stop asking me.” (RS34)*
>
> *“[…] since we last spoke, I have had a few additional lines to submit to hPSCreg, but these were required to go through the full submission and approval process, although were approved within 10 min so not sure how thoroughly the data gets assessed.” (RS38)*
>
> *“But again, I think that some people, some researchers might not be, you know, entirely honest with their cell lines, and they just need, they just want the name out there. So they just, you know, put up a karyotype from when it was originally done, you know, five years ago or something and then, or bounced through like five students hands or something and you don’t know what it actually is now. So that made me feel not really sure if it was that useful in that sense. (RS34)”*

Other negative experiences are associated with the design of the registry, with two users revealed concerns about the time effort required to register all their cell lines in the future, and one user complained about the difficulty in searching on hPSCreg based on characteristics of a line. Additionally, one user expressed frustrations about hPSCreg’s use of institutional administrators (RS21). The system automatically designates the first registrant for an institution as the “administrator registrant,” granting them the authority to approve or deny subsequent user registrations from the same institution (Figure 1), regardless of whether they hold an administrative role within the institution.

> *“A bit painful. In fact, yeah, it was really, it was labour-intensive. I had to actually get [Institute’s name] Intellectual Property and Commercialisation manager [name] to contact the registry and have the [Institute’s name] put on as a host institute.” (RS21)*

## 4. Discussion

In summary, our research results suggest that most Australian researchers have limited awareness of international stem cell registries and are unclear about their purposes. The few participants who have used hPSCreg described their primary motivation for registration was to meet publication prerequisites, with the idea of enhancing the visibility of their cells typically being a secondary consideration. The lack of understanding regarding the FAIRness of cell lines and their associated data in the stem cell field, coupled with the unclear goal of registration and misunderstandings about certain hPSCreg processes, was associated with a series of negative experiences reflected in the interviews. Ultimately, this has impacted users’ perceptions of the platform’s utility and the reliability of its data.

Unlike the EC in the European Union, which requires its funding recipients to register cell lines used in funded research projects, and obtain a certificate from hPSCreg attesting to their ethical provenance and biological properties, Australia lacks a funding requirement to register cell lines with hPSCreg. Therefore, the primary incentive for Australians to register with hPSCreg stems from publication requirements, which also serves as the main avenue through which Australians learn about hPSCreg. Without uptake of ISSCR registration recommendations by Australian funders, there seems to be little external impetus to register cell lines by Australian researchers.

While journal *Stem Cell Research* contributes to a certain level of Australian hPSC registration, at least half of our interviewees treated registration as a hurdle to publication rather than a resource. Consequently, Australian researchers were rarely engaging with the primary goals of the registry. First, as a centralized information hub, its function for disseminating information is not widely adopted among Australian researchers. Secondly, regarding information collection, even well-intentioned researchers may not provide all the data needed for full registration in the rush to publication, consequently most of Australian lines on hPSCreg remain partially registered. Thirdly, most of the researchers that we interviewed were unaware of the purpose and value of registration, although two groups did adopt hPSCreg nomenclature or documentation without formally registering their lines. Manual adoption of naming without registration poses the potential risk of duplicating names identical names of different lines might be generated through hPSCreg, resulting in cell line identity confusion, and exacerbating issues in linking data with associated cell lines, contradicting the purpose of registration.

Partially registered cell lines with standard UI do provide some level of public visibility of cell lines and do enable information sharing across data resources such as BioSample, Cellosaurus, and Wikidata. However, superficial information also impacts on the usefulness of the platform. For researchers searching for a cell line, the first step involves assessing whether a cell line is suitable for their research. This entails understanding details about how the line was made, disease groups associated with the cell line, and any restrictions on cell line availability because of the type of donor consent obtained for the line. Explicit ethical provenance, including understanding what a donor has consented to is crucial in ensuring the ethical conduct of future research. Most researchers would be aware of instances where a historical lack of consent around cell lines has led to accusations of exploitation, exemplified by the HeLa cell tragedy (Masters, 2002). Moreover, researchers need to know whether a cell line derived from a healthy donor, has undergone genetic correction, or carries a particular disease of interest. The absence of disease-related details poses a challenge for researchers in their decision-making. For registrants who didn’t complete the registration, their cell lines will have limited visibility on the platform since they are hard to retrieve by simple characteristic-related keyword search besides its UI. The awareness of the existence of incomplete information in hPSCreg, as exemplified by RS34, also impacts on how much trust the user places in the data entered from other registrants. From a registry perspective, this indicates that superficial engagement, usually due to forced registration, has unintended consequences that may lead to poor user perceptions in the platform.

The misunderstandings that we identified in our Australian user group, especially about what information hPSCreg collects and how hPSCreg validates this data, underscores a deeper issue about user engagement. For instance, misunderstandings about whether identifiable information was collected in the informed consent and which data field is mandatory or optional arise primarily due to registrants’ carelessness, given the instructions on hPSCreg documentation and prompts beside these fields. However, frustrations also arose from user difficulties navigating hPSCreg requirement for a designated institutional authority. The perceptions reported here about the platform’s user-friendliness and ease of registration may be biased by the small sample size and should not be considered representative of all hPSCreg users.

It is worth noting that after the Journal *Stem Cell Research, Stem Cell Reports*, the Journal of the ISSCR, is now asking authors to complete a checklist that asks for a UI from a registry. This checklist is being trialled between October 2023 and March 2024 before the editorial board decides on whether to make the checklist a publication hurdle (NIH hESC Registry, 2023). It seems likely that, in the future, there will be more mandatory requirements for registration. To foster better registration practices among researchers, we propose the importance of raising awareness about the current data-linking issues and highlighting the purpose and value of registration. We also encourage active communication between registries and their users to ensure easy-to-use platforms that are fit for purpose.

### Limitations

This qualitative investigation into Australian researchers’ experiences and attitudes towards hPSCreg involved a small sample of individuals who have used hPSCreg, contextualised within an Australian setting.

## Conclusion

Our study provides the first systematic data regarding hPSCreg users’ experience since its establishment in 2007. Overall, our study found that mandatory registration through publications serves as a double-edged sword—while it ensures a certain level of registration, it also gives rise to many issues rooted in the stem cell community that haven’t fully embraced the importance and value of registration. As ISSCR guidelines recommend registration, we anticipate a broader advocacy for registration in various forms, including partnership with Journals and stem cell related initiatives. We propose to actively communicate registration goals and value to the stem cell community. Additionally, we propose the establishment or improvement of an easy-to-use registry that fits to purpose.

## Authorship contributions

Conceptualization; MH, CAW

Data curation; DS, MH, NM

Formal analysis; MH

Funding acquisition; RA, DN, CAW

Investigation; MH, DS, EL

Project administration; RA, CAW

Supervision; RA, CAW

Roles/Writing - original draft; MH

and Writing - review & editing, MH, DS, EL, DN, AK, NM, SM, RA, CAW

## Acknowledgments

The authors acknowledge discussions with the EAOR team.

**Appendix A. Supplementary data**

## Supplementary Data 1

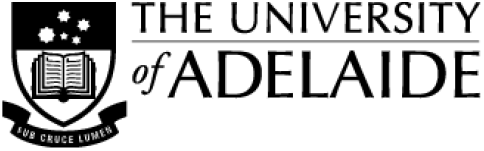

### Human Research Ethics Committee (HREC)

#### CONSENT FORM (interviews and observations)

1. I have read the attached Information Sheet and agree to take part in the following research project:

2. I have had the project, so far as it affects me, and the potential risks and burdens fully explained to my satisfaction by the research worker. I have had the opportunity to ask any questions I may have about the project and my participation. My consent is given freely.
3. Although I understand the purpose of the research project, it has also been explained that my involvement may not be of any benefit to me.
4. I agree to participate in the activities outlined in the participant information sheet. Participating in an interview is required to participate in the Delphi study.
  a. Interview ⍰ Yes ⍰ No
  b. Observation ⍰ Yes ⍰ No ⍰ N/A
  c. I would be willing to be contacted about participation in the Delphi Study in the future: ⍰ Yes ⍰ No
5. I agree to be:
  a. Audio recorded ⍰ Yes ⍰ No
  b. Video recorded ⍰ Yes ⍰ No
  I agree to have:
  a. Photos of my work taken (not identifying people or the laboratory) ⍰ Yes ⍰ No ⍰ N/A
  b. Anonymised data relating to my interview/observations shared via Figshare ⍰ Yes ⍰ No
6. I understand that I am free to withdraw from the project, where applicable: 1) before data analysis starts, two months after the interview takes place, and/or 2) before the analysis starts, two months after the completion of the Delphi study (i.e., after the final workshop).
7. I have been informed that the information gained throughout the project may be published in journal articles, conference presentations, a website, news articles and press releases, and a report to the funding body.
8. I have been informed that while I will not be named in the published materials, it may not be possible to guarantee my anonymity given the nature of the study and/or small number of participants involved. I understand that I will be given the opportunity to review any direct quotes that will be used in publications as detailed in the Participant Information Sheet.
9. I agree to my information being used for future research purposes as follows: Research undertaken by **these same researcher(s)**
  a. on an extension of, or a project closely related to, the current project: ⍰ Yes ⍰ No
  b. in the same general area of research: ⍰ Yes ⍰ No
  Research undertaken by **any researcher(s)**
  a. on an extension of, of a project closely related to, the current project: ⍰ Yes ⍰ No
  b. in the same general area of research: ⍰ Yes ⍰ No
10. I understand my information will only be disclosed according to the consent provided, except where disclosure is required by law.
11. I am aware that I should keep a copy of this Consent Form, when completed, and the attached Information Sheet.

|  |  |
| --- | --- |
| **Title:** | **Developing an Evidence-Based Model for Building Trust in Australian Stem Cell Research and Therapies** |
| **Ethics Approval Number:** | **H-2022-095** |

#### Participant to complete

Name:_______________________ Signature: _______________________ Date: _______________________

#### Researcher/Witness to complete

I have described the nature of the research to_______________________

*(print name of participant)*

and in my opinion she/he understood the explanation.

Signature: _______________________ Position: _______________________ Date: _______________________

## Supplementary Data 2

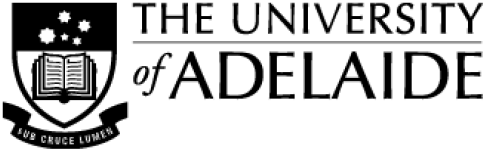

### PARTICIPANT INFORMATION SHEET

**PROJECT TITLE:** Developing an Evidence-Based Model for Building Trust in Australian Stem Cell Research and Therapies

**HUMAN RESEARCH ETHICS COMMITTEE APPROVAL NUMBER:** H-2022-095

**PRINCIPAL INVESTIGATOR:** Professor Rachel Ankeny

Dear Participant,

You are invited to participate in the research project described below.

#### What is the project about?

This research project focuses on testing a model for building more transparency and trust in medical research/therapeutics related to the use of stem cells. In recent years, there have been efforts to ensure that the data and results of research are openly shared and include the perspectives of all stakeholders who will be affected by the research results. We intend to address a significant gap in the evidence base for improving community trust, acceptance, and adoption of stem cell-based therapies in Australia by using a ‘commons’ model. In a commons model of governance, the responsibility of governing a common good rests with all those who are to share the benefits provided by these goods.

#### Who is undertaking the project?

This project is being conducted by Professor Rachel Ankeny, Professor Dianne Nicol, Professor Joan Leach, Professor Christine Wells, Dr Dan Santos, Dr Edilene Lopes McInnes, Ms Rebekah Harms, and Ms Mengqi Hu.

#### Why am I being invited to participate?

You are being invited because of your professional background and experience in stem cell research, which fits the selection criteria of our research project. We are interested in hearing from a range of stakeholders about their views on stem cell research and therapies in Australia.

We will also conduct another study (using the Delphi method) to assess the commons model that we will develop based on the interviews and other research. We may invite you again to take part in this other study if you indicate that you are willing to participate in future. The Delphi study will be conducted through an online process. We anticipate that we will have around three workshops (around 1.5 hours each) and two rounds of completing surveys on your own (around 30 minutes each) on non-consecutive days.

You can choose if you want to take part only in the interview and/or observation (if relevant), and whether you wish to be re-contacted about the later Delphi study. We will provide you with a consent form where you will be able to select your preferences in terms of which parts of the project you wish to participate in.

#### What am I being invited to do?

You are being invited to take part in a semi-structured interview that will last up to 60 minutes. The interview can be conducted face-to-face at your workplace, in a public space convenient for you, or via an online platform. We will need to audio-record (if face-to-face) or video-record (if online, but we will only use the audio of the interview for data analysis) the interview to produce a transcript that will be analysed. Before the interview, we will ask you to fill out a short survey (around 5-10 minutes) to let us know about your work and some demographic information (e.g., gender, age group) so we can contextualise your answers when doing the analysis.

For a small number of researchers, we also wish to do a brief observational study, which will last approximately 60 minutes. We will come to the laboratory to observe researchers whilst conducting their work in the laboratory to understand their work practices associated with stem cells. We may also ask some questions to understand what is being done and why. Researchers may take down notes of activities observed, and request to audio-record your explanations and responses to questions and take photos of lab activities (with no people or identifying features of the lab).

Later we may invite you to take part in a Delphi study where we will ask participants to answer questions and to rate, in order of relevance, statements related to a commons model for stem cell research in Australia. This study is conducted online, through surveys (completed individually) and workshops (completed in a group). The group workshops will be recorded to produce a transcript. You will be provided more information and be asked to consent, if you choose to participate in this particular part of the project.

#### How much time will my involvement in the project take?

The interview will take approximately 60 minutes and the observation an additional 60 minutes if relevant. The later Delphi study will occur online, with participation not to exceed six hours over three non-consecutive days.

#### Are there any risks associated with participating in this project?

We do not anticipate that the research will cause risks to participants. You may feel uncomfortable discussing some topics around the applications of stem cell research and therapies. If this occurs, we will pause the interview. You do not have to answer a question if it makes you feel uncomfortable and you have the right to withdraw at any time and this will not affect you negatively in any way. With respect to observation, participants may experience inconvenience whilst being observed carrying out experimental laboratory work. Questions may be asked during this lab work, but the topics of these questions and discussions will only relate to the work being conducted – no personal, non-project related questions or topics will be asked, and participants are free to decline to answer any question for whatever reason should they wish.

#### What are the potential benefits of the research project?

There is no immediate benefit to you personally. But we hope this project will help to build a model of governance (commons framework) for stem cell research in Australia that could potentially increase access and equity to future stem cell therapies for a diverse population, and build trust and transparency in stem cell research and therapeutics.

#### Can I withdraw from the project?

Participation in this project is completely voluntary. If you agree to participate, you can withdraw from the study two months after the interview and/or observation (before we start the data analysis, when it becomes difficult to separate the codes generated from the interview). In the case of the Delphi study, you may withdraw from the study up to two months after the completion of the final workshop.

#### What will happen to my information?

When we contact you, we will keep your personal information (e.g., full name, email address, telephone number, workplace) in the University electronic systems, where it is protected by password or, if in hard copy, in a locked cabinet at the school of Humanities at the University of Adelaide. Only research members will have access to this information, which will be destroyed after five years.

For the data analysis phase of the study, all transcripts will be de-identified and coded. Transcription companies will receive de-identified information and keep the data in similar conditions to the University.

At the end of the project, we intend to convene four half-day workshops involving a range of stakeholders to present our final framework. We also plan to submit at least three peer-reviewed journal articles and promote the framework through a website. We will define how to promote the framework to community organisations in consultation with them. In all of these events and products, we intend to keep the participants’ privacy by taking reasonable steps to not identify them (e.g., using codes and not naming any entities cited during the interview). However, due to the small field of stem cell research in Australia, some participants (particularly researchers and patient organisation representatives) may be identifiable despite these measures.

If we want to use one of your quotes in any work, we will allow you to see part of the draft publications and give you two weeks to respond if you are happy for us to include this extract. If you do not respond within this timeframe, we will understand that as an agreement to use the quote. We will try to contact you in various ways during these two weeks. If you are interested, we can offer you a copy of this project’s final report.

In the accompanying consent form, we will ask for your consent to use this data in future research related to stem cell research/therapies as a way of allowing us and other researchers to explore this topic in more depth.

Your information will only be used as described in this participant information sheet, and it will only be disclosed according to the consent provided, except as required by law.

#### Who do I contact if I have questions about the project?

If you have any questions about this project, you can primarily contact Prof Rachel A. Ankeny -, + 61 8 8313 5570. You can also contact:

Dr Edilene Lopes McInnes -, +61 8

8313 0617 Dr Dan Santos -

Ms Rebekah Harms -

Ms Mengqi

(Chi-Chi) Hu -

Prof Dianne Nicol -, +61 3 6226 7553

Prof Joan Leach -, +61 2 6125 4513

Prof Christine Wells -, + 61 3 8344 3795

#### What if I have a complaint or any concerns?

The study has been approved by the Human Research Ethics Committee at the University of Adelaide (approval number H-2022-095). This research project will be conducted according to the *NHMRC National Statement on Ethical Conduct in Human Research 2007 (Updated 2018*). If you have questions or problems associated with the practical aspects of your participation in the project, or wish to raise a concern or complaint about the project, then you should consult the Principal Investigator. If you wish to speak with an independent person regarding concerns or a complaint, the University’s policy on research involving human participants, or your rights as a participant, please contact the Human Research Ethics Committee’s Secretariat on:

Phone: +61 8 8313 6028

Post: Level 3, Rundle Mall Plaza, 50 Rundle Mall, ADELAIDE SA 5000

Any complaint or concern will be treated in confidence and fully investigated. You will be informed of the outcome.

#### If I want to participate, what do I do?

If you decide to participate in our project, all you have to do is to contact us, and we will be in touch with you to organise your participation (including asking your consent to participate formally in the research). You can contact Dr Dan Santos, or Ms Rebekah Harms.

Yours sincerely,

Prof Rachel Ankeny, Prof Dianne Nicol, Prof Joan Leach, Prof Christine Wells, Dr Edilene Lopes McInnes, Dr Dan Santos, Ms Rebekah Harms, and Ms Mengqi (Chi-Chi) Hu.

## Supplementary Data 3

Pre-interview survey

_______________________________________________________________________

**Start of Block: Consent**

CONSTENT

**PRE-INTERVIEW SURVEY**

***Informed Consent* Survey Investigators:** Dr Dan Santos, Ms Mengqi (Chi-Chi) Hu

**Project title:** Developing an Evidence-Based Model for Building Trust in Australian Stem Cell Research and Therapies

**Institutions:** Australian National University, University of Melbourne

You are invited to participate in a short survey about your experiences with and opinions of stem cell registries. Demographic and research related information will also be collected. You may skip questions you don’t want to answer. Please complete and submit the survey before your scheduled interview.

If you have any questions about this survey, please contact Ms Mengqi (Chi-Chi) Hu or Dr Dan Santos.

Please refer to the Participant Information Sheet that was provided earlier for more information about the research project, including additional contact information if you have further questions.

∘ Yes I consent (1)
∘ No I do not consent (2)

*Skip To: End of Block If PRE-INTERVIEW SURVEY Informed Consent Survey Investigators: Dr Dan Santos, Ms Mengqi (Chi-Chi) Hu*… *= Yes I consent*

*Skip To: End of Survey If PRE-INTERVIEW SURVEY Informed Consent Survey Investigators: Dr Dan Santos, Ms Mengqi (Chi-Chi) Hu*… *= No I do not consent*

**End of Block: Consent**

_______________________________________________________________________

**Start of Block: Demographics**

Q1 We require your name and institution to help us ask tailored questions for a post survey interview. This information will be kept strictly confidential.

Gender, age range, and role are collected to understand your personal experience of registries.

-----------------------------------------------------------------------------------------------------------

Q1 What is your name?

________________________________________________________________

-----------------------------------------------------------------------------------------------------------

Q2 What is your institution?

________________________________________________________________

________________________________________________________________

________________________________________________________________

________________________________________________________________

________________________________________________________________

-----------------------------------------------------------------------------------------------------------

Q2 What is your gender?

∘ Male (1)
∘ Female (2)
∘ Non-binary / third gender (3)
∘ Prefer not to say (4)

-----------------------------------------------------------------------------------------------------------

Q3 What is your age?

∘ 18-24 years (1)
∘ 25-34 years (2)
∘ 35-44 years (3)
∘ 45-54 years (4)
∘ 55-64 years (5)
∘ 65 + years (6)

-----------------------------------------------------------------------------------------------------------

Q4 How do you obtain hPSC lines? (you may select more than one options)

□ I make my own lines (1)
□ I obtain from others (2)
□ other ways (6) __________________________________________________

**End of Block: Demographics**

_______________________________________________________________________

**Start of Block: Manufacturers**

*Display This Question:*

*If How do you obtain hPSC lines? (you may select more than one options) = I make my own lines*

Q5a We would like to hear more about your work.

-----------------------------------------------------------------------------------------------------------

*Display This Question:*

*If How do you obtain hPSC lines? (you may select more than one options) = I make my own lines*

Q4a Approximately, how many pluripotent stem cell lines have you made?

________________________________________________________________

-----------------------------------------------------------------------------------------------------------

*Display This Question:*

*If How do you obtain hPSC lines? (you may select more than one options) = I make my own lines*

Q4b Have your pluripotent stem cell lines been used outside your lab?

∘ Yes (1)
∘ No (2)

*Display This Question:*

*If How do you obtain hPSC lines? (you may select more than one options) = I make my own lines And Have your pluripotent stem cell lines been used outside your lab? = Yes*

Q4c Do you know where they were used?

∘ Yes (1)
∘ No (2)

*Display This Question:*

*If How do you obtain hPSC lines? (you may select more than one options) = I make my own lines*

*And Do you know where they were used? = Yes*

*And Have your pluripotent stem cell lines been used outside your lab? = Yes*

Q4d Could you share any related information as examples, such as publications related to the use of these pluripotent stem cell lines? (For example, please describe or enter DOIs)

__________________________________________________________________________

**End of Block: Manufacturers**

_______________________________________________________________________

**Start of Block: used a registry?**

Q5 Have you used a stem cell registry?

∘ Yes (1)
∘ No (2)

*Display This Question:*

*If Have you used a stem cell registry? = Yes*

Q5a Which stem cell registry/registries have you used?

□ hPSCreg (1)
□ SKIP (2)
□ Other (please specify) (3) ________________________________________________

**End of Block: used a registry?**

_______________________________________________________________________

**Start of Block: purpose of using a registry**

*Display This Question:*

*If Have you used a stem cell registry? = Yes*

Q5b What did you use a registry for?

________________________________________________________________

________________________________________________________________

________________________________________________________________

________________________________________________________________

__________________________________________________________________________

**End of Block: purpose of using a registry**

_______________________________________________________________________

**Start of Block: Manufacturers with stem cell registries**

*Display This Question:*

*If How do you obtain hPSC lines? (you may select more than one options) = I make my own lines And Have you used a stem cell registry? = Yes*

\ We would like to hear more about your experience with stem cell registries.

-----------------------------------------------------------------------------------------------------------

*Display This Question:*

*If How do you obtain hPSC lines? (you may select more than one options) = I make my own lines And Have you used a stem cell registry? = Yes*

Q4c How many of your pluripotent stem cell lines are registered in **${lm://Field/1}?**

________________________________________________________________

________________________________________________________________

________________________________________________________________

________________________________________________________________

________________________________________________________________

**End of Block: Manufacturers with stem cell registries**

_______________________________________________________________________

**Start of Block: evaluate existing regisry**

*Display This Question:*

*If Have you used a stem cell registry? = Yes*

Q5c Do you think **the existing registries** are useful to you and people in your field?

∘ Strongly Agree (1)
∘ Agree (4)
∘ Neutral/ Unsure (5)
∘ Disagree (6)
∘ Strongly Disagree (7)

**End of Block: evaluate existing regisry**

_______________________________________________________________________

**Start of Block: Prospect of ASCreg**

*Display This Question:*

*If Have you used a stem cell registry? = No*

Intro to ASCretg A **pluripotent stem cell registry** is a publicly accessible and registerable online database that centralises stem cell metadata. Specifically, a registry allows registrants to upload their stem cell metadata and enables users to search for cell lines in the database. It is distinguished from a stem cell bank or a biobank where the actual cell lines are generated, stored and managed.

The metadata stored on a pluripotent stem cell registry may include donor attributes, ethics, derivation, culture methods, characteristics, relevant publications, etc. Access to a physical cell line requires private consultation through the contact of the registrant.

-----------------------------------------------------------------------------------------------------------

Q6 If an **Australian Pluripotent Stem Cell Registry** was established to capture relevant information about stem cells generated in Australia, do you think it would be useful to you and people in your field?

∘ Strongly Agree (1)
∘ Agree (2)
∘ Neutral/ Unsure (3)
∘ Disagree (4)
∘ Strongly Disagree (5)

**End of Block: Prospect of ASCreg**

_______________________________________________________________________

**Start of Block: score through the benefit list**

*Display This Question:*

*If Have you used a stem cell registry? = Yes*

Q39 We would like to understand your opinions of **existing stem cell registries**.

Please score as follows:

1. Strongly Disagree
2. Disagree
3. Neutral/ Unsure
4. Agree
5. Strongly Agree

-----------------------------------------------------------------------------------------------------------

*Display This Question:*

*If Have you used a stem cell registry? = Yes*

Q35 **Do you think that the existing registries are helpful for**

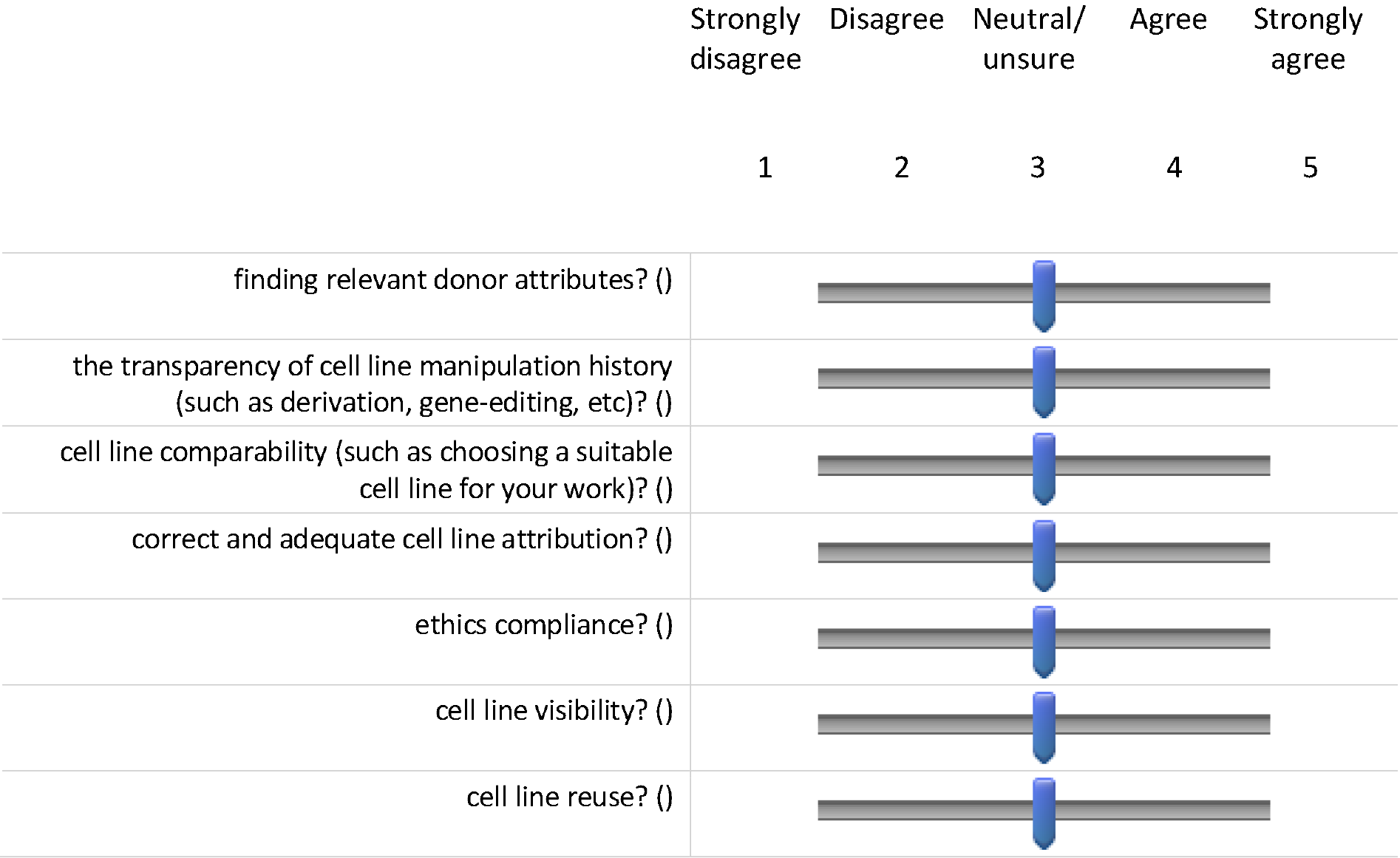

*Display This Question:*

*If Have you used a stem cell registry? = Yes*

Q36 **Do you think that the existing registries serve as platforms for**

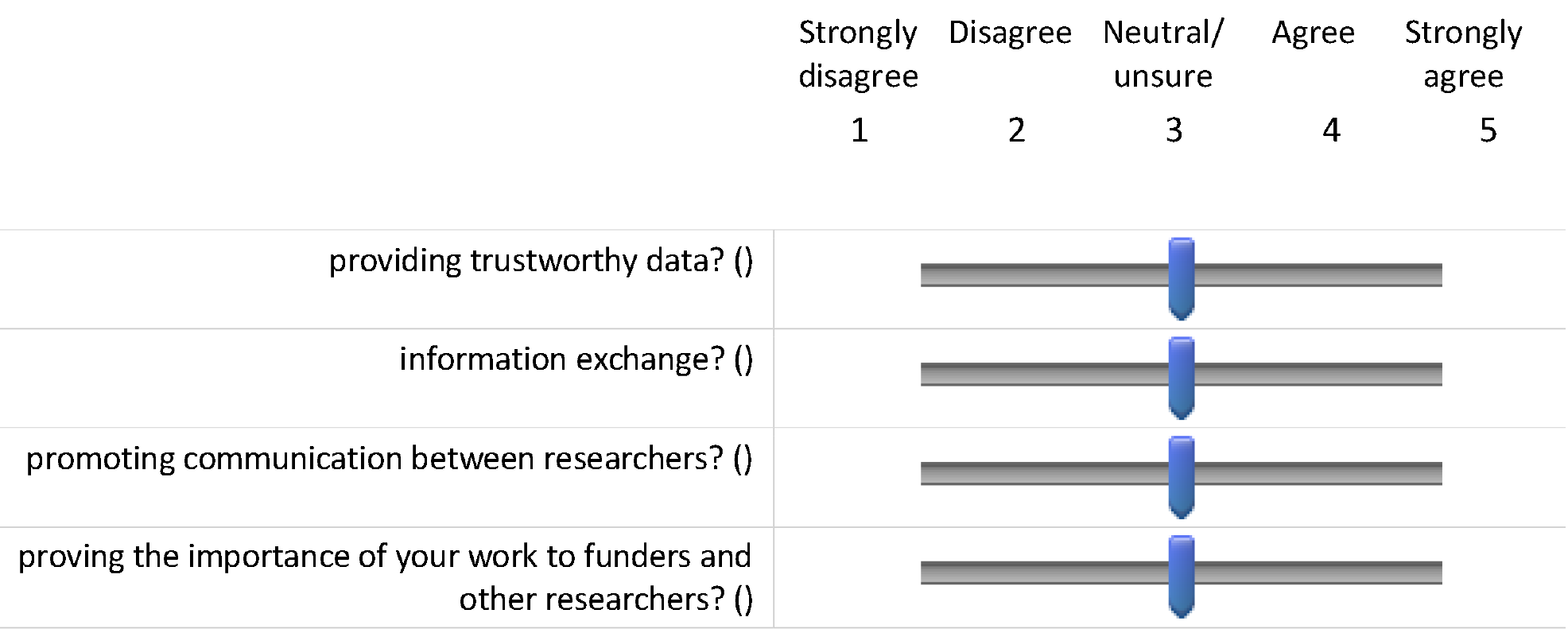

**End of Block: score through the benefit list**

_____________________________________________________________

